# Alcohols inhibit glucose transporters noncompetitively at a constant membrane concentration according to lipid theory

**DOI:** 10.1101/2025.11.20.689647

**Authors:** Yukifumi Uesono

## Abstract

Various chemicals obey the Meyer-Overton correlation, in which potency increases exponentially with the partition coefficient. However, the principle remains unclear because of controversy between the lipid and protein theories. As the protein theory is based on the competitive inhibition of the non-membrane protein luciferase by alcohols and anesthetics, kinetic analysis was conducted on yeast hexose transporter Hxt2, a membrane protein. *n*-Alcohols (C_2_-C_8_) inhibited Hxt2 noncompetitively and drug efflux according to the correlation. Thus, the alcohols interacting nonspecifically with membrane affect various membrane targets and inhibit Hxt2 from the lateral membrane-facing side, thereby supporting the lipid theory. The alcohols exerted similar potencies at a constant membrane concentration (*C*_m_), regardless of the chain length. Consequently, the principle of the Meyer-Overton correlation is the maintenance of a constant *C*_m_, whereas administered aqueous concentrations (*C*_w_) only decrease with their partition coefficients. Intracellular *C*_m_, but not extracellular *C*_w_, provides an accurate measure of chemical potency.

## INTRODUCTION

Lipophilic drugs exert their effects at low doses and are considered potent. This is because many of these drugs obey the Meyer-Overton (MO) correlation, in which potency increases exponentially with lipophilicity.^1,2^ The MO correlation is observed in various biological effects, such as anesthesia, toxicity, antimicrobial activity, hemolysis of red blood cells, and plant cell growth, induced by numerous chemicals, including alcohols, alkanes, fatty acids, antipsychotic drugs, and agrochemicals.^3-13^ Therefore, the MO correlation represents a fundamental principle describing chemicals with organism interactions and has led to the development of quantitative structure−activity relationship (QSAR) models used to estimate biological activity from chemical structures.^11-13^ However, the potencies of chemicals are typically evaluated based on the administered concentration in aqueous solution (*C*_w_), which may not accurately reflect the true potency of lipophilic chemicals governed by the MO correlation.

Alcohols exhibit diverse biological effects, including antimicrobial, toxic, and anesthetic activities. The minimum effective concentration (MEC) decreases exponentially with increasing lipophilicity (chain length), thus obeying the MO correlation. Consequently, alcohols have historically been studied as model compounds to understand the principle of MO correlation. The MO correlation led to the concept of lipid membrane; based on this, the lipid theory was proposed, stating that chemicals, including alcohols, dissolve directly into the lipophilic membrane and modulate various membrane targets indirectly.^4^ However, highly lipophilic long-chain alcohols (≥C_13_) lose their biological activity despite being predicted to exhibit strong effects according to the lipid theory.^3,14^ The lipid theory failed to explain this cutoff paradox, instead, the protein theory was proposed.^15^ The protein theory is based on the competitive inhibition of firefly luciferase (FFL) by alcohols and anesthetics in the absence of lipid membranes.^16^ The theory advocates that the alcohols bind proteins directly and explains the cutoff by assuming the ideal pockets accommodating alcohols with various chains, except for long-chain alcohols. However, FFL is not a membrane protein, and the pocket has not been identified. Consequently, the principle underlying the MO correlation has not been elucidated and remains controversial.

The cutoff paradox of the long-chain alcohols is attributed to their low saturated aqueous solubility (*S*_w_), which is the maximum concentration of an effective hydrated monomer capable of dissolving in the membrane, thereby decreasing the membrane concentration (*C*_m_).^17^ Because of the insufficient *C*_m_, they do not reach the MEC required in the membrane and therefore lose their biological activity. Increasing *C*_m_ can restore strong activity by allowing MEC to be reached, thereby avoiding the cutoff and adhering the MO correlation.^17,18^ In addition, the half inhibitory concentrations (IC_50_) for the domain formation of giant unilamellar vesicles (GUVs) decrease exponentially with increasing chain lengths, indicating that the MO correlation is established within the membrane itself.^19^ These observations collectively support the lipid theory as a reasonable explanation for the MO correlation. To determine which theory is correct, it is necessary to determine how alcohols act on the practical membrane proteins rather than on model proteins that lack membrane context. This study describes the inhibition mechanisms of nonionic alcohols on the yeast hexose transporter Hxt2, the differences with cationic amphiphilic drugs (CADs), the principle of the MO correlation, and a method for evaluating chemical potency according to the lipid theory.

## RESULTS

### *n*-Alcohols inhibit hexose transporter function according to the MO correlation

In budding yeast, *n*-alcohols (C_1_-C_3_) induce glucose starvation and inhibit growth,^8^ similar to the effects of CADs such as local anesthetics, antipsychotic phenothiazines (chlorpromazine: CPZ),^20^ and antimalarial drugs (quinacrine: QC).^21^ CPZ and QC localize to the yeast cell membrane and noncompetitively inhibit the function of the hexose transporter (Hxt),^21^ suggesting that various alcohols, despite being nonionic, inhibit Hxt function. Yeast Hxts belong to the major facilitator superfamily^22^ and are structurally and functionally conserved in the human glucose transporters GLUT1 and 4.^23^ These transporters specifically recognize monosaccharides and facilitate their uptake by cells through passive transport driven by alternating conformational changes.^22^ Because Hxts are encoded by 18 genes whose expression is tightly regulated by extracellular hexose concentrations,^24^ kinetic analysis in wild-type cells is challenging; the coexistence of multiple isozymes with different *K*_m_ values makes it difficult to determine accurate kinetic parameters. Moreover, Michaelis–Menten equation analysis requires measurement of the initial rate of glucose import in a very short time without a contribution from export. Failure to meet these conditions can lead to incorrect *K*_m_ determination and misinterpretation such as allosteric inhibition. To overcome these challenges, the HXT2m strain was used, in which 18 *HXT* genes were deleted. This strain expresses only the high-affinity glucose transporter *HXT2* and was used to evaluate the effect of alcohols on the initial rate of 2-deoxyglucose (2DG) uptake (zero-trans influx) using a bioluminescence method.^21^

Alcohols (C_2_-C_8)_ inhibited the initial rate of 2DG uptake in a dose-dependent manner (Figure 1A), and their half-inhibitory concentrations (Hxt2-IC_50_) decreased exponentially with increasing chain length (Figure 1B). In MO correlations, lipophilicity was quantitatively expressed as partition coefficients: the octanol/water partition (*P*_ow_: *C*_o_/*C*_w_) or lipid membrane/water partition (*K*_mw_: *C*_m_/*C*_w_). The logarithmic IC_50_ values were linearly correlated with the logarithms of *K*_mw_ for alcohols, with a slope of approximately -1 (Figure 1C). These results indicate that alcohols target Hxts as membrane proteins and inhibit their functions in accordance with the MO correlation.

**Figure 1.**
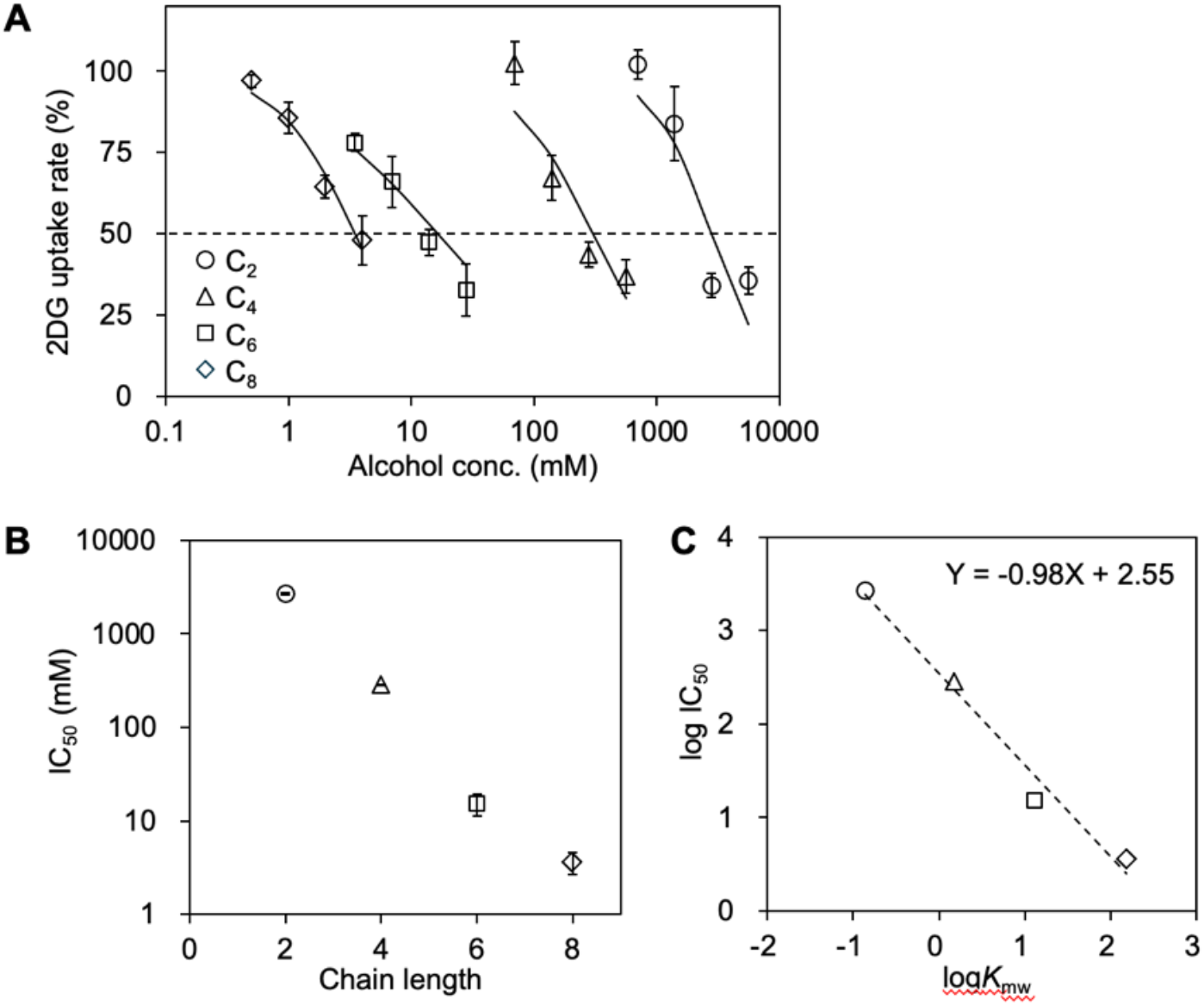
Alcohols inhibit the function of hexose transporter according to the Meyer–Overton (MO) correlation. (A) Initial rates of 2-deoxyglucose (2DG) uptake were measured in Hxt2m yeast cells treated with alcohols (C_2_–C_8_) for 30 min. Rates in alcohols-treated cells are indicated as percentages of those in untreated cells and plotted against the administered aqueous concentrations of alcohols (*C*_w_). Logistic curves are fitted to determine the half-maximal inhibitory concentrations (IC_50_) for the alcohols and indicated as averages. (B) IC_50_ values of the alcohols are plotted against their chain length. Data are the means (SE) (*n* ≥3). (C) IC_50_ values of the alcohols are plotted against the logarithms of membrane-water partition coefficient (*K*_mw_) (Table 1). The equation of linear regression was calculated using Excel and shown on the right side.

### *n*-Alcohols noncompetitively inhibit hexose transport by Hxt2 irrespective of chain length

The alcohol-induced inhibition of the membrane domain formation of GUVs may be attributable to alterations in the composition of lipids forming the domains caused by the nonspecific penetration of alcohols into membrane lipids.^19^ The IC_50_ values for the inhibition (GUV-IC_50_) were highly correlated with the minimum inhibitory concentrations (MIC) for yeast growth^17^ and half effective dose (ED_50_) for tadpole anesthesia,^25,19^ although their specific membrane targets are unknown. Because the GUV-IC_50_ values were highly correlated with the Hxt2-IC_50_ values (Figure 2), alcohols may inhibit Hxt2 function from the lateral side of the membrane, consistent with the lipid theory. However, alcohols may directly inhibit Hxt2 function, as observed for luciferase inhibition, which supports the protein theory. To distinguish between these theories, kinetic analysis of Hxt2 in the presence of a biomembrane is necessary (Figure 3).

**Table 1.**
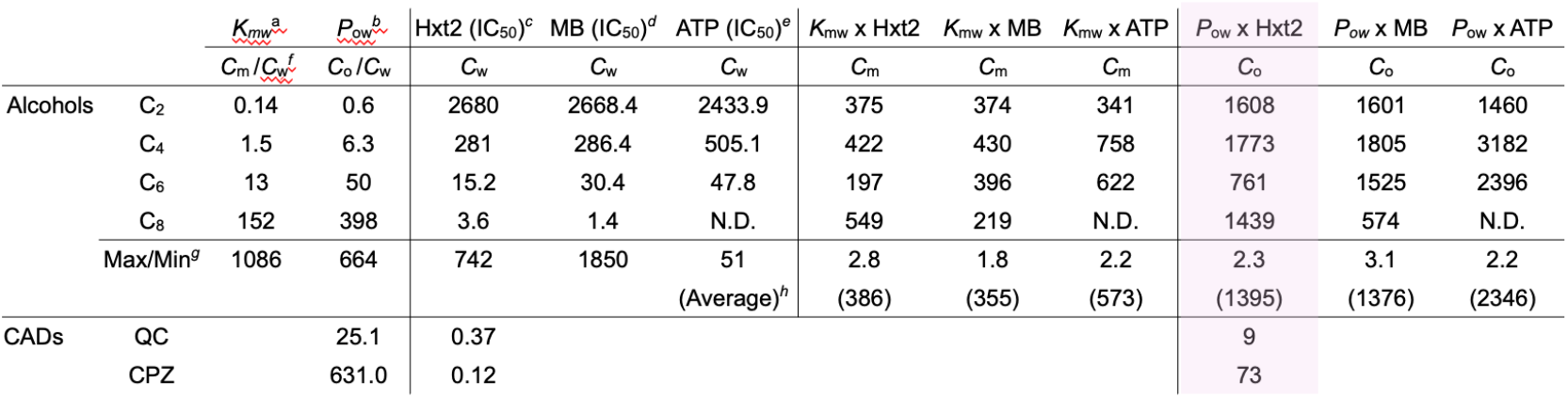
^*a*^Membrane−water partition coefficients^3^ and ^*b*^octanol−water partition coefficients of alcohols were calculated from log*D* at pH 7.4 estimated using MarvinSketch software (2025, Oct.). ^*c*^IC_50_s of Hxt2 function, ^*d*^IC_50_s of methylene blue (MB) efflux, and ^*e*^IC_50_s of intracellular ATP level were determined in Figure 1 and 4, respectively, under the same conditions as used for the HXT2m strain. ^*f*^Concentration (*C*: mM) in different phases are expressed as *C*_m_ (in membrane), *C*_w_ (in water), and *C*_o_ (in octanol). ^*g*^Ratios of the maximum and minimum values in each column (C_2_-C_8_). ^*h*^Averages of *C*_m_ and *C*_o_ among alcohols are shown in parentheses.

**Figure 2.**
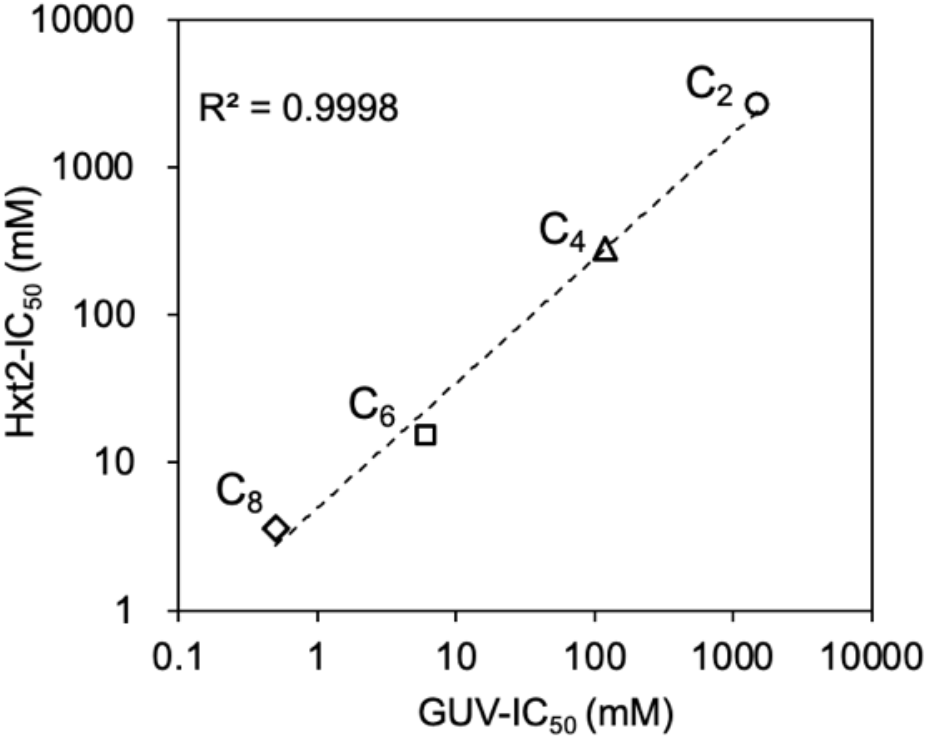
Inhibitions of domain formation in giant unilamellar vesicles (GUVs) reflect inhibition of hexose transporter. Half-maximal inhibitory concentrations (IC_50_) values of the alcohols for the initial rate of 2-deoxyglucose (2DG) uptake by Hxt2 are plotted against the IC_50_ values of GUV domain formation.^19^ The coefficients of determination between two variables were calculated using Excel and are shown on the left sides.

**Figure 3.**
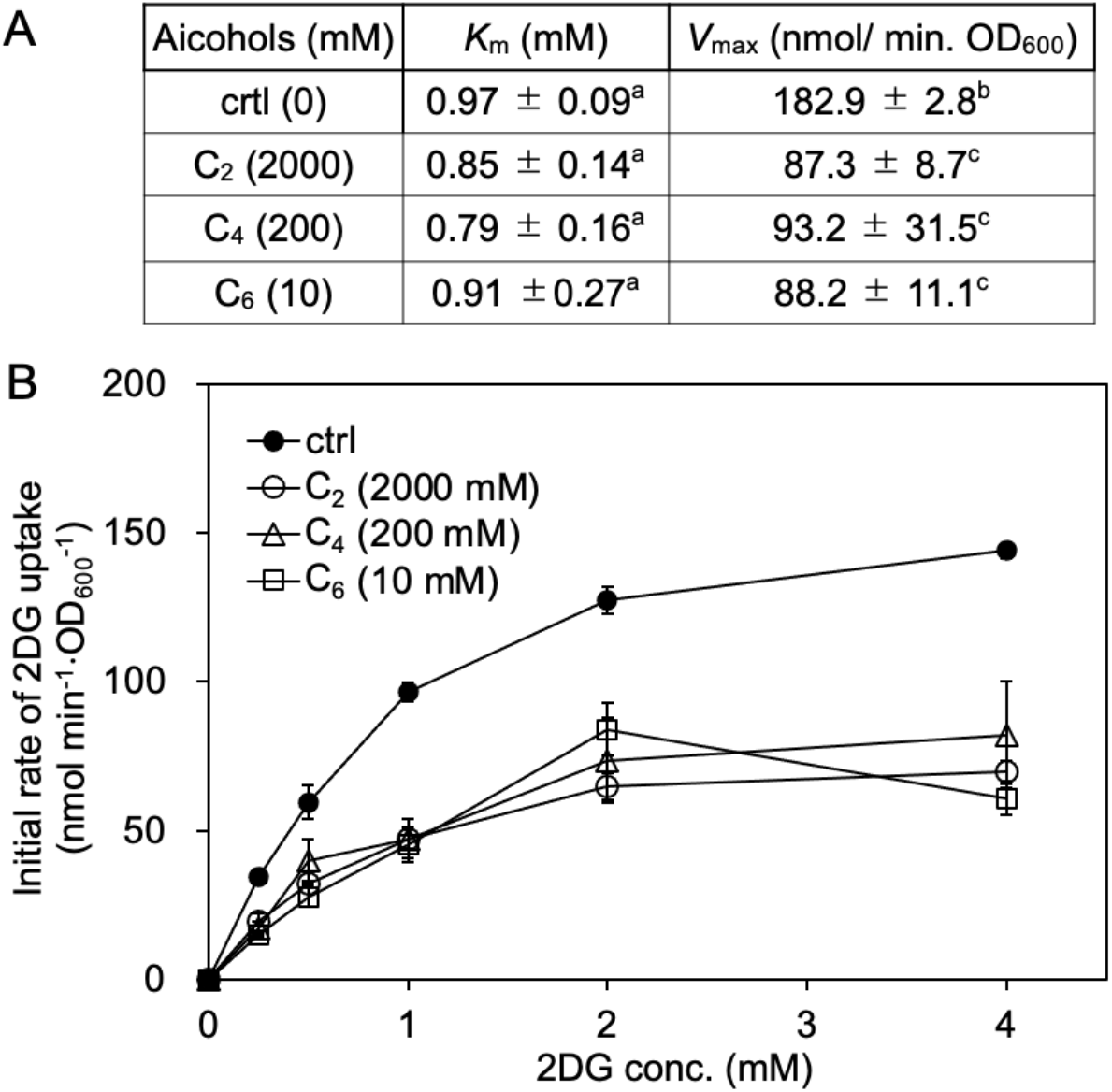
Alcohols inhibit Hxt2 noncompetitively irrespective of chain length. (A) Kinetic parameters for 2-deoxyglucose (2DG) uptake inhibition by alcohols at pH 7.4. Data are the means (SE) (*n* ≥3). Statistical significance was determined using Tukey−Kramer multiple-comparison test. Different letters denote significant differences (p < 0.05). (B) Initial rate of 2DG uptake by Hxt2 in the presence of alcohols approximately their half-maximal inhibitory concentration (IC_50_) values are plotted against the 2DG concentrations.

The kinetic parameters for alcohol-induced inhibition of 2DG uptake were determined in the HXT2m strain. In the absence of alcohols, the *K*_m_ of Hxt2 was approximately 0.8–0.9 mM, similar to but slightly lower than values previously reported (1.5 and 3.1 mM) using a radioisotope uptake assay in an *hxt*1-7 null mutant expressing *HXT2*.^26,27^ The *V*_max_ (624 nmol min^-1^ [mg of cells]^-1^, equivalent to 183 nmol min^-1^ OD_600-1_) was higher than that reported previously (97 nmol min^-1^ [mg of cells]^-1^),^26^ possibly because of the high expression of *HXT2* from a multicopy vector. Therefore, the bioluminescence method is available for kinetic analysis instead of the radiolabeling method. In the presence of alcohols (C_2_, C_4_, and C_6_) near their IC_50_ values, *V*_max_ decreased by approximately half, whereas *K*_m_ did not change (Figure 3A). The saturation curves for the 2DG uptake rate versus the increase in 2DG displayed typical Michaelis–Menten behavior rather than sigmoid kinetics (Figure 3B). These results indicate that alcohols noncompetitively inhibit 2DG transport via Hxt2, independent of chain length.

### *n*-Alcohols inhibit drug efflux activity according to the MO correlation

Because alcohols nonspecifically penetrate membranes, they may affect a broad range of membrane proteins other than Hxts. Viable cells exhibit drug efflux activity, whereas the dead cells do not. When yeast cells are mixed with a solution containing methylene blue dye (MB), the MB concentration in the solution decreases as the number of dead cells increase, as MB accumulates in dead cells. Thus, the MB decreases in solution reflects the decreases in drug efflux activity and cell viability. Spectroscopic measurement of the MB concentration provides a simple method to evaluate the effects of alcohols on these cellular functions.^28^

The HXT2m strain was treated with alcohol for 30 min, and then the cell pellets were mixed with MB solution. Alcohols (C_2_-C_8_) decreased the MB concentration in the supernatant in a dose-dependent manner, indicating inhibition of drug efflux activity and reduced cell viability (Figure 4A). The IC_50_ values (MB-IC_50_) for MB efflux decreased exponentially with increasing chain length, indicating that inhibition of drug efflux activity by alcohols obeyed the MO correlation (Figure 4B). The MB-IC_50_ values for each chain length were similar to the Hxt2-IC_50_ values for 2DG transport inhibition (Figure 4B).

**Figure 4.**
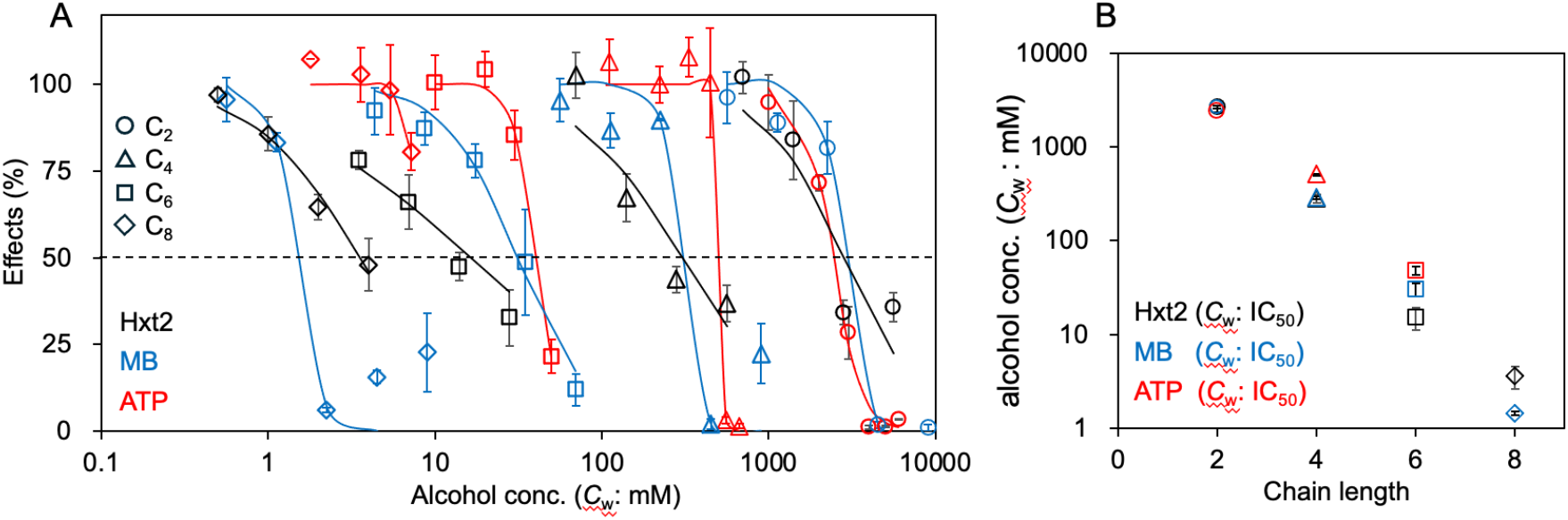
Alcohols decrease drug efflux activity and intracellular ATP levels, according to the Meyer–Overton (MO) correlation. The efflux activities of methylene blue (MB) and intracellular ATP levels were measured in HXT2m cells treated with alcohols (C_2_–C_8_) for 30 min. MB levels in the supernatant (blue shapes) and ATP levels (red shapes) of the alcohol-treated cells are indicated as percentages of those of untreated cells and plotted against the administered aqueous concentrations of alcohols (*C*_w_). For comparison, the percentages of 2-deoxyglucose (2DG) uptake rates in Figure 1 are indicated (black shapes). These logistic curves are fitted to determine their half-inhibitory concentrations (IC_50_) for the alcohols and are indicated by the averages. (B) IC_50_ values for these effects judged by *C*_w_ are plotted against their chain length. Data are the means (SE) (*n* ≥ 3).

Thus, decreases in drug efflux activity and viability occurred at concentrations close to those required for inhibition of 2DG transport. In the dose-response curves, MB efflux activity was inhibited by alcohols (C_2_-C_6_) at slightly higher concentrations than those required to inhibit 2DG transport, except for C_8_ (Figure 4A, black and blue lines). Because glucose starvation alone does not decrease cell viability in budding yeast,^29^ these results indicate that alcohols impair additional membrane-associated targets during inhibition of glucose transport, thereby supporting nonspecific interactions of alcohols with the lipid membrane.

### n-Alcohols decrease intracellular ATP level according to the MO correlation

In the presence of glucose, budding yeast generate ATP via the glycolytic pathway with suppressing the respiratory pathway. As glucose deprivation rapidly and transiently decreases intracellular ATP levels in yeast,^29^ alcohol-induced inhibition of glucose uptake may decrease ATP levels. The HXT2m strain was treated with alcohols (C_2_-C_8_) for 30 min in YPD, a yeast complete medium containing 2% glucose, and intracellular ATP levels were measured. Alcohols C_2_-C_6_ decreased intracellular ATP levels in a dose-dependent manner, whereas C_8_ showed only partial inhibition (Figure 4A). The IC_50_ values for ATP levels (ATP-IC_50_) decreased exponentially with increasing chain length (C_2_-C_6_), indicating that alcohol-induced ATP depletion obeyed the MO correlation (Figure 4B). The ATP-IC_50_ value for C_2_ was similar to the Hxt2-IC_50_ and MB-IC_50_ values, consistent with their dose-response curves (Figure 4A and B), indicating that C_2_ induced these effects at similar concentrations. In contrast, the ATP IC_50_ values for C_4_ and C_6_ were higher than their respective Hxt2-IC_50_ and MB-IC_50_ values (Figure 4A). The ATP IC_50_ value for C_8_ could not be determined because of only partial inhibition, even at high concentrations (Figure 4A, red line); thus, C_8_ is a cutoff compound for decreased ATP under these conditions, possibly because of insufficient cellular accumulation of C_8_ within 30 min.^18^ These results indicate that the alcohols (C_4_-C_8_) require higher concentrations to reduce ATP levels than to inhibit glucose uptake or drug efflux. Therefore, Hxt2 transport and drug efflux are likely both directly inhibited by alcohols, rather than indirectly inhibited by decreased ATP levels.

### Alcohols exert similar effects at a constant membrane concentration regardless of chain length

According to protein theory, the potency of alcohols for the above biological effects was evaluated based on their administered *C*_w_. However, when alcohols nonspecifically penetrate the membrane and act on various membrane targets, as proposed by lipid theory, the *C*_m_, rather than *C*_w_, must be evaluated because the *C*_m_ around the membrane target determines actual potency.

In aqueous solutions containing cells surrounded by membrane, chemicals partition into the membrane and water according to *K*_mw_, enabling estimation of the *C*_m_ values of alcohols as the product of *K*_mw_ (*C*_m_/*C*_w_) with their administered *C*_w_ values.^3,18,19^ Using this approach, the *C*_m_ values (*C*_2_–*C*_8_) calculated as the product of *K*_mw_ with Hxt2-IC_50_ (*C*_w_) differed by 2.8-fold, whereas that among Hxt2-IC_50_s was 742-fold. Similar trends were observed when *C*_m_ was calculated from MB-IC_50_ and ATP-IC_50_ respectively (Table 1). These results indicate that alcohols exert similar potency after reaching a constant *C*_m_ irrespective of chain length, although the administered *C*_w_ decreases exponentially with chain length.

The constant *C*_m_ required for ATP reduction was slightly higher than that needed for Hxt2 and MB efflux inhibition (Table 1). To determine the detailed differences in *C*_m_ for each effect, the dose-response curves for *C*_m_ were compared. C_2_ induced these effects with similar *C*_m_ values (Figure 5). In contrast, the *C*_m_ values for C_4_ and C_6_ increased in the following order: Hxt2 inhibition < MB efflux inhibition < ATP reduction. The *C*_m_ values of C_8_ increased in the following order: MB efflux inhibition < Hxt2 inhibition < ATP level decrease, which differed from the order of C_4_ and C_6_. These results indicate that alcohols act on similar membrane targets and alcohols (C_4_-C_8_), but not (C_2_), affect additional membrane targets as *C*_m_ increases.

**Figure 5.**
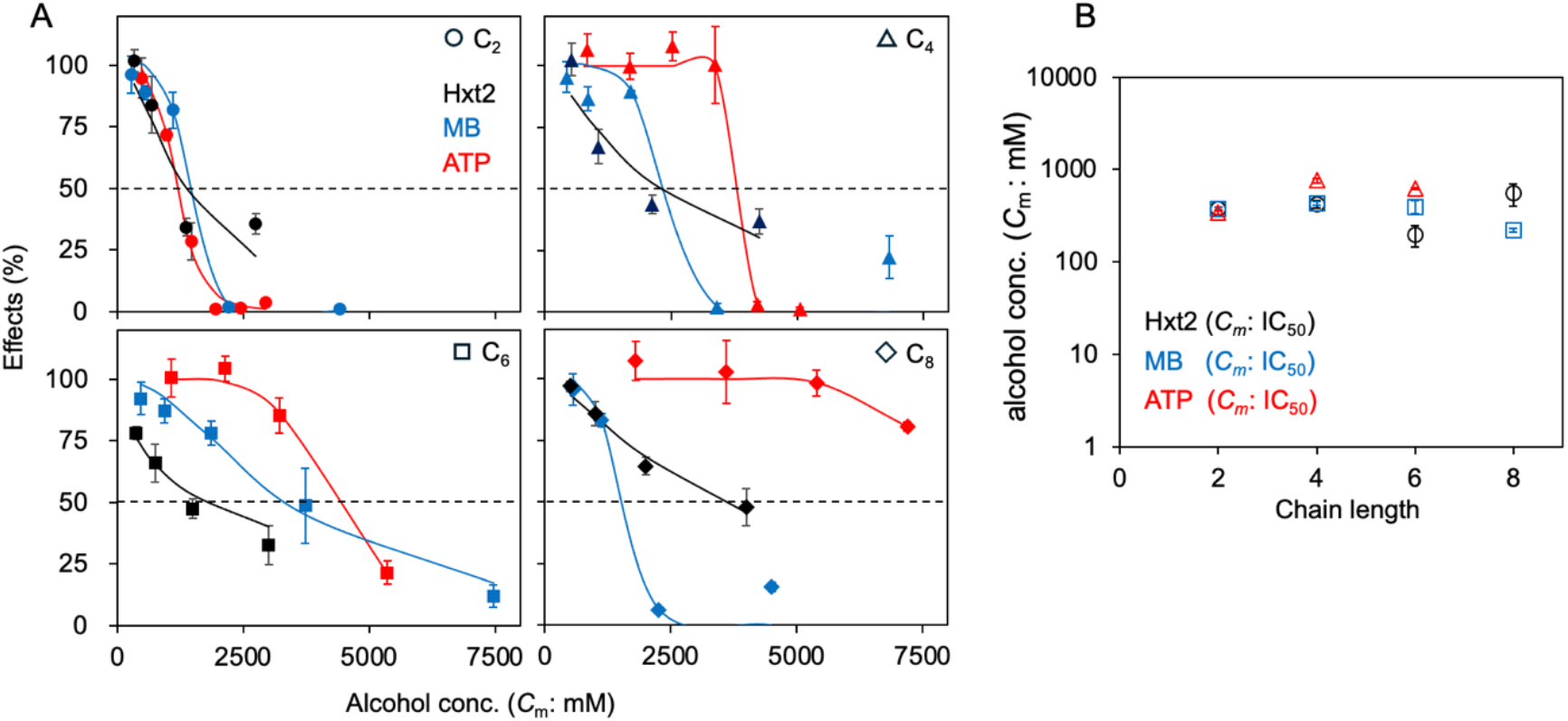
Alcohols exert biological effects at similar membrane concentrations. (A) Percentages for 2-deoxyglucose (2DG) uptake rates, methylene blue (MB) efflux activities, and intracellular ATP levels in Figure 4 are plotted against the membrane concentrations of each alcohol (*C*_m_), respectively. *C*_m_ values were calculated as the product of the administered concentration in water (*C*_w_) and the membrane-water partition coefficients (*K*_mw_: *C*_m_/*C*_w_) of each alcohol.^3^ The shapes and logistic curves are same as those in Figure 4, respectively. (B) IC_50_ values for these effects judged by *C*_m_ are plotted against their chain length.

When an alcohol does not reach a certain *C*_m_, it does not exert the effect, as predicted by the cutoff paradox of long-chain alcohols (>C_12_).^14,3^ The prediction has been experimentally validated to exert the effects by increasing the *C*_m_ of the cutoff alcohols using methods for microorganisms such as increasing the *S*_w_ of alcohols by melting at high temperature, using mutants defective in the drug efflux pump,^17^ and prolonging the exposure time to cells.^18^ Because *C*_m_ is defined as the amount of chemicals per unit membrane, decreasing the amount of membrane may increase the relative *C*_m_. When the number of cells, and thus the total membrane, decreased, C_14_ inhibited yeast growth, thereby escaping the cutoff (Figure 6A). These findings indicate that a certain *C*_m_ for each biological effect must be reached before alcohols exert their effects. Collectively, alcohols act on similar membrane targets at the constant *C*_m_ values, respectively, and that the targets increase as the *C*_m_ increases, although their detailed mechanisms appear to vary with chain length.

**Figure 6.**
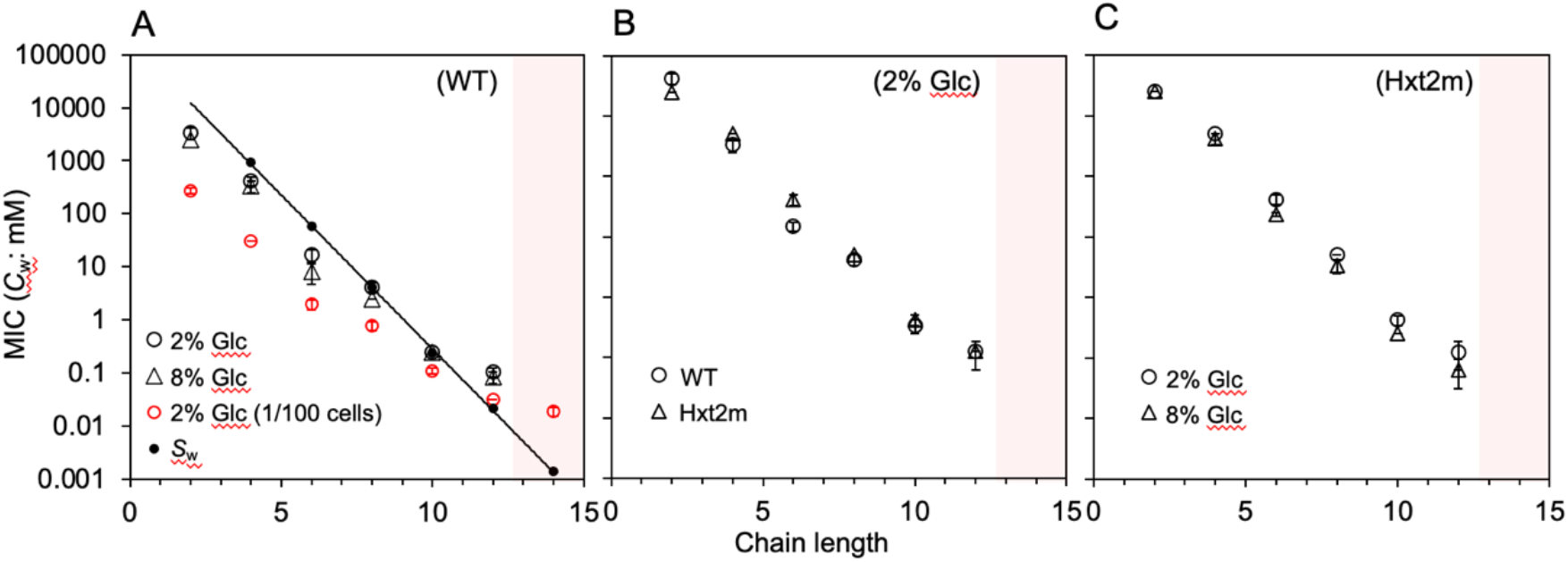
Effects of cell number and glucose concentration on the yeast minimum inhibitory concentration (MIC) of alcohols. (A) BY4741 strains (wild-type: 6.5 × 10^4^ cells) were incubated statically with alcohols (C_2_-C_14_) for 3 days at 25 °C in YPD containing 2% (circle) and 8% glucose (triangle). The diluted BY4741 strains (1/100: 6.5 × 10^2^ cells) were incubated in YPD containing 2% glucose under the above condition (red circle). Saturated water solubilities (*S*_w_) of the alcohols are indicated as black filled circles and a line. These MIC and *S*_w_ values are plotted against the chain length. The red zone contains the cutoff alcohols (C_14_) with no growth inhibitory effect. (B) BY4741 and HXT2m strains (6.5 × 10^4^ cells) were incubated statically with alcohols in YPD containing 2% glucose at 25 °C for 3 days (circle) and 5 days (triangle), respectively, and their MIC values were determined. (C) HXT2m strains (6.5 × 10^4^ cells) were incubated statically with alcohols for 5 days at 25 °C in YPD containing 2% (circle) and 8% glucose (triangle). Data are the means (SE) (*n* ≥3).

### Alcohols inhibit glucose transport at membrane concentrations higher than CADs

In budding yeast, various CADs inhibit glucose uptake and growth, thereby inducing glucose starvation state.^20^ Additionally, CADs, such as CPZ and QC, inhibit Hxt2 function noncompetitively.^21^ The resulting CAD-induced growth inhibition can be partially rescued by excess glucose, indicating that impaired hexose transport causes growth inhibition.^20,21^ In contrast, alcohol-induced growth inhibition was not alleviated by excess glucose in the wild-type and HXT2m strains (Figure 6A and C). To determine the differences in the inhibition mechanisms of CADs and alcohols, their *C*_m_ values were compared. However, the actual *K*_mw_ values for most compounds are unknown, and determining these values for electrolytes such as QC and CPZ is difficult because their partition coefficients vary with pH-induced structural changes.^21^ MarvinSketch software for quantitative structure−property relationship (QSPR) can estimate pH-dependent partition coefficient (*P*_ow_: *C*_o_/*C*_w_) from the structure as log*P* or log*D*.^30^ Because Pow approximates *K*_mw_, the corresponding *C*_o_ values were used as surrogates for *C*_m_ to compare the membrane concentrations among CADs and alcohols.

Although the *C*_o_ values (761–1773 mM) calculated from Hxt2-IC_50_ (*C*_w_) for C_2_–C_8_ were larger than their *C*_m_ values, the difference was 2.3-fold, which was similar to 2.8-fold among their *C*_m_ values (Table 1, *C*_o_). Therefore, *C*_o_ can be used to evaluate *C*_m_. Because *P*_ow_ for CADs varies with pH, whereas those for nonionic alcohols do not, the *C*_o_ values of CADs estimated using the *P*_ow_ at pH 7.5 were compared with those of alcohols. The average *C*_o_ for C_2_–C_8_ was 1395 mM, whereas those for QC and CPZ were 9 and 73 mM, respectively (Table 1, *C*_o_). This result indicates that alcohols require *C*_m_ values of 150-fold for QC and 19-fold for CPZ to inhibit glucose transport with similar potency. Thus, the high *C*_m_ for Hxt2 inhibition may explain why alcohol-induced growth inhibition cannot be alleviated by excessive glucose.

## DISCUSSION

### Alcohols inhibit Hxt2 function from the lateral side of the membrane

This is the first report to explain the MO correlation by focusing on the kinetic mechanism of an actual membrane protein, the glucose transporter. A central question is how alcohols inhibit membrane hexose transporters. In GUVs composed of cholesterol and saturated and unsaturated phospholipids, the lipids spontaneously associate with each other and form a biased distribution, the liquid-ordered (L_o_) and disordered (L_d_) domains.^31^ The L_o_ domain containing saturated lipids is a physicochemical basis for lipid raft accumulating various receptors and channels in biomembranes.^32^ Alcohols disrupt domain formation in protein-free GUVs by dispersing the biased lipid distribution through nonspecific penetration into the membrane, an effect similar to that of excessive cholesterol.^19^ Yeast Hxts, however, are uniformly distributed in the plasma membrane and do not localize to lipid rafts.^20,33^ Thus, Hxt2 inhibition is likely to directly arise from alcohols dissolving nonspecifically into the membrane surrounding Hxt2 but not the secondary effects by raft disruption. Consistent with this, Hxt2 showed noncompetitive inhibition by alcohols (Figure 3), indicating that alcohols act at sites other than the hexose recognition site. Taken together, these findings suggest that alcohols inhibit Hxt2 function from the lateral membrane side adjacent to the transporter.

### Principle of MO correlation is to maintain membrane concentration of chemicals constant to exert the same effect

The IC_50_ values as *C*_w_ for Hxt2 function, MB efflux, and ATP level decreased exponentially with increasing alcohol chain length (partition coefficients) (Figure 4); however, their IC_50_ values as *C*_m_ were nearly constant, irrespective of their chain length (Table 1, Figure 6B). These results indicate that chemicals exert similar biological effects when they reach the same *C*_m_, as observed in anesthesia of tadpole,^3,14,34^ inhibition of microbial growth,^18^ and domain formation in artificial membranes,^19^ which is the principle of the MO correlation. *C*_w_ only decreases exponentially according to the partition coefficients to maintain the same *C*_m_. The concentration reflecting the actual potency is *C*_m_, which was taken into cells, but not *C*_w_, which could not be taken into cells.

At the molecular level, a constant *C*_m_ reflects the contact frequency in the membrane between the alcohol and Hxt2 in the membrane rather than changes in membrane properties. When this frequency reaches a certain threshold in the membrane, regardless of the chain length, structural changes in Hxt2 are inhibited from the lateral side, resulting in the noncompetitive inhibition of glucose transport (Figures 3,7). The EC_50_s for anesthesia of tadpole^3^ and MICs for yeast growth were highly correlated with the GUV-IC_50_s for domain formation of artificial membranes,^19^ as observed for the Hxt2-IC_50_s (Figure 2). As these *C*_m_ values are constant, regardless of the chain length, their mechanisms may be similar to those of Hxt2 inhibition based on nonspecific penetration into the membrane, although their membrane targets remain unknown.

**Figure 7.**
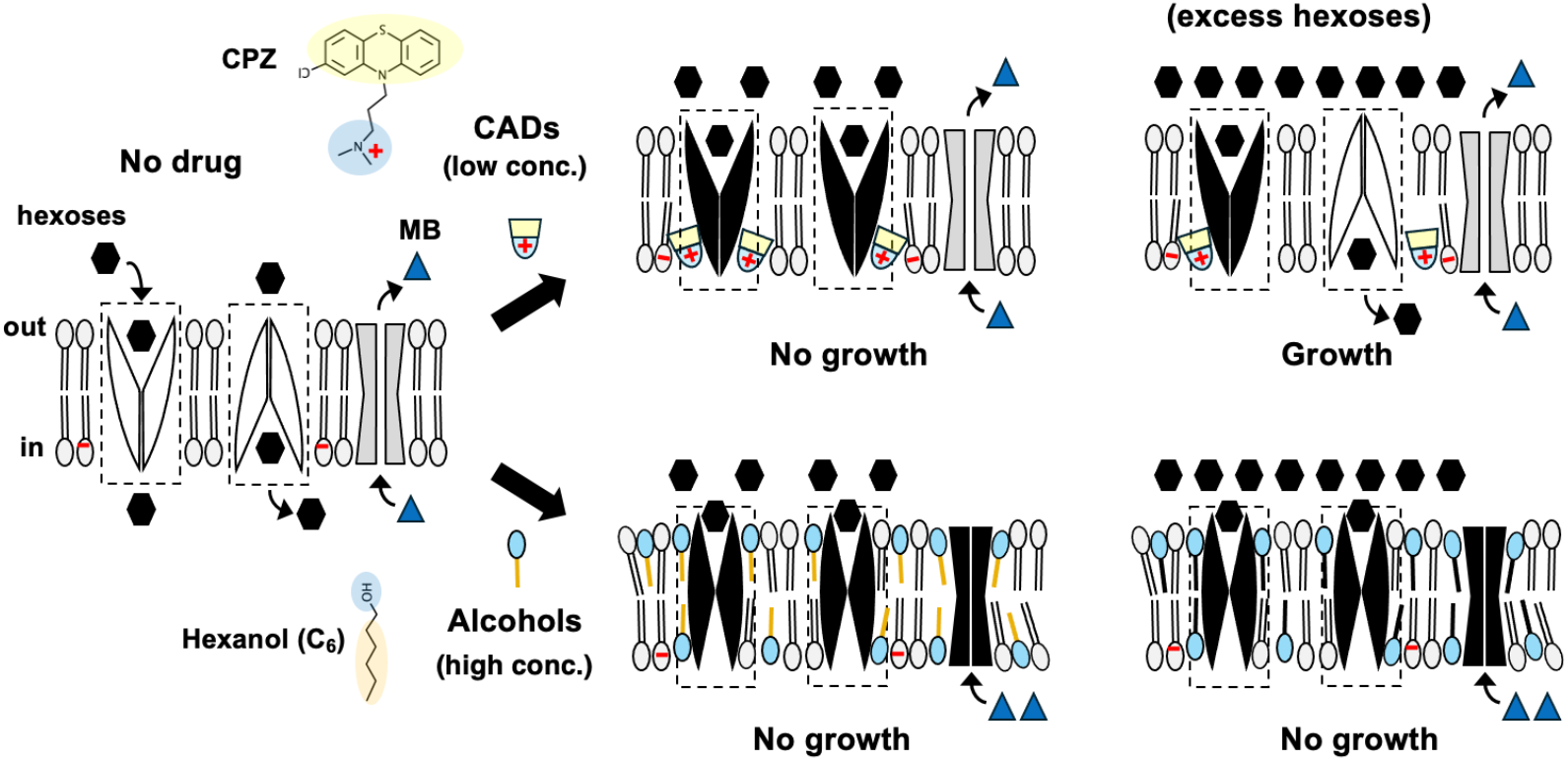
Mechanisms of inhibition of hexose transporter by alcohols and cationic amphiphilic drugs (CADs). Hexoses are selectively transported across membranes via oscillated conformational changes of Hxts. The oscillation areas are surrounded by dotted boxes. Although the pump for methylene blue (MB) efflux is unidentified, hexoses and methylene blue are indicated as hexagonal and triangular shapes, respectively (left). CADs such as chlorpromazine (CPZ) and quinacrine (QC) penetrate nonspecifically into the negatively charged inner leaflets of lipid membrane along the amphiphilic orientation. Because the hydrophobic region of CAD broadens laterally in the leaflets, even when small amounts enter the oscillation area, they prevent conformational changes to the lateral side, thereby inhibiting glucose uptake (upper center, black shapes). However, excessive glucose activates oscillation via efficient interactions of Hxts with hexoses and prevents CADs from penetrating the oscillating area, thereby alleviating CAD-induced growth inhibition and conferring resistance (upper right, white shape). In contrast, nonionic alcohols penetrate nonspecifically into both leaflets of lipid membrane because of no charges. Because the straight alkyl chains of alcohols fit with the alkyl chains of the leaflets, large amounts of alcohols are required to suppress the structural change of Hxts by compressing the entire structure from the lateral side of both leaflets. Thus, this suppression affects various membrane proteins, thereby inhibiting glucose uptake and drug efflux (lower center, black shapes). The suppression to the entire structure compress Hxts even when the oscillation is already active; thus, alcohol-induced growth inhibition is not alleviated by excess glucose (lower right, black shapes).

Increasing the *C*_m_ of alcohols (C_4_-C_8_) increased their biological effects (Figure 5A). The local anesthetic tetracaine, a CAD, also inhibits glucose uptake at low concentrations and additively phosphorylates eIF2 alpha in the amino acid starvation pathway at high concentrations.^35^ Because these chemicals dissolve in the membrane nonspecifically via their amphiphilic structure, increasing *C*_m_ increases the number of membrane targets, regardless of their structural difference, thereby producing additive or off-target effects. At more higher concentrations, nonspecific penetration of these chemicals into membrane destroys membrane structure and lead to cell lysis as their toxicities.^8,20,21^ These observations support lipid theory.

### Alcohols and CADs inhibit Hxt function through different mechanisms

The inhibition of yeast growth by CADs, such as local anesthetics (TC, lidocaine, and dibucaine), antipsychotic phenothiazines (CPZ and trifluoperazine),^20^ and antimalarial QC,^21^ was alleviated by excess glucose, regardless of structural differences, whereas that caused by alcohols was not (Figure 6A, C). These differences reflect distinct inhibitory mechanisms. Hexoses are selectively transported across membranes via oscillating conformational changes in Hxts that typically have outward-facing structures to recognize external hexoses in yeast.^20,22^ CADs nonspecifically penetrate the membrane based on the orientation of amphiphilic phospholipids. Because the heads of the inner leaflets of the plasma membrane are enriched in negatively charged phospholipids,^36^ the positively charged hydrophilic region of CADs interact electrostatically with the inner leaflets,^20,21^ and the hydrophobic region spreads laterally at the hydrophobic region of the inner leaflets. When a small amount of CAD enters the oscillation area for Hxts, their structural changes to the inward-facing form are prevented, thereby inhibiting hexose transport (Figure 7, middle upper, black shapes). However, when the oscillation is already active by excessive hexose, the CADs could not enter the oscillation area, thereby allowing transport and alleviating growth inhibition (Figure 7, right upper, white shapes), which is a novel mechanism for drug resistance as reported previously.^20^

In contrast, nonionic alcohols penetrate lipid bilayers leaflets along their amphiphilic orientations. Because the alkyl chains of alcohols do not spread to the lateral side of the membrane, it is difficult to prevent structural changes in Hxts by entering the oscillation area in small amounts, as observed for CADs. Thus, alcohols may suppress the structural change in Hxts by pressing the entire structure from the lateral side of both leaflets; however, in this case, large amounts of alcohol would be required in the membrane (Figure 7, middle lower, black shapes). This mechanism is consistent with the much higher *C*_o_ values of alcohols compared with those of CADs (Table 1), supporting the lateral pressure mechanism^37^. Because this suppression by high *C*_m_ affects the Hxt conformation globally, it can press even the oscillated transporter activated by excess glucose (Figure 7, right lower, black shapes). It differs from the mechanism that prevents a change in the inward-face sructure of the inner leaflets using small amounts of CADs. Thus, this mechanism accounts for lack of excess glucose-mediated rescue of alcohol-induced growth inhibition (Figure 6A, C) and induction of various effects (Figure 4).

Various amphiphilic compounds, such as alcohols and CADs, would contact with the transmembrane domains of Hxt2 at the protein-lipid interface as a secondary effect of nonspecific interactions with the membrane, rather than via specific binding. Therefore, if structural analyses of membrane proteins identify contact sites in transmembrane domains^38^ only with membranes but not without membranes, such findings would support lipid theory rather than protein theory.

### Competitive inhibition of FFL by alcohols is not a basis for protein theory

These mechanisms, based on the noncompetitive inhibition of Hxt, clearly differ from the protein theory. The protein theory explains direct interactions between proteins and chemicals via competitive inhibition of FFL with alcohols in the absence of a membrane,^39^ and the cutoff paradox of long-chain alcohols for anesthesia by assuming ideal pockets accommodating various chains but not long-chains.^16^ FFL was found to be a family of long-chain acyl-CoA synthetase (ACSL) that adds AMP to fatty acids.^40^ Because the luciferase activity of ACSL^41^ and long-chain fatty acyl-AMP formation activity of FFL have been confirmed,^42^ FFL is a special enzyme with a high affinity for alkyl chains. Structural analysis of ACSL proposed a narrow tunnel structure in which the alkyl chains enter, with an active center at the back.^43^ Competitive inhibition would be reasonable for alcohols and luciferin entering this tunnel.^39^ However, the tunnel is not the ideal pocket because FFL efficiently forms acyl-adenylates from long-chain fatty acids (>C_18_) as a substrate,^42^ contradicting the protein theory explained by assumption of chain length constraints. Indeed, this tunnel has not been found in GABA or NMDA receptors with no luciferase or ACSL activity, which are membrane-target candidates for anesthetics and alcohol. In addition, the cutoff paradox of long-chain alcohols is due to their low *C*_m_ values^18^ and can be avoided by increasing the *C*_m_ using a mutant defective in the drug efflux pump,^17^ prolonging exposure to cells^18^, and reducing the cell membrane (Figure 6A). Accordingly, an ideal pocket in the protein is not essentially required to explain this paradox. Despite FFL is not membrane protein, presumably, overinterpretation extending the case of special enzyme to membrane proteins led to serious confusion: protein theory.

### Cutoff compounds with long alkyl chains may contribute to diabetes

Because general anesthetics decrease glucose uptake in the human brain,^44^ glucose starvation in the brain may be involved in loss of consciousness, similar to hypoglycemic comas in patients with diabetes.^45^ CADs such as local anesthetics and antipsychotic phenothiazines induce surgical diabetes.^46,47^ Ethanol induces insulin-resistant hyperglycemia in rodents,^48,49^ suggesting that alcohol inhibits glucose transporters such as GLUT1 expressed throughout the body, including in the brain, rather than insulin-dependent GLUT4. Because Hxts are structurally and functionally conserved from yeast to humans,^23^ these amphiphiles, regardless of their structural difference, may inhibit glucose transport into human tissues in a manner similar to that in yeast (Figure 7), resulting in hyperglycemia.

Studies should focus on cutoff alcohols with long alkyl chains. Their high *P* values reduce the *S*_w_ and limit the *C*_m_ to below the MEC in the membrane.^18^ Thus, early inhibition of glucose transport would not be detected, and these compounds would appear to be nontoxic. However, over time, the cutoff alcohols accumulate with slow degrees in membranes during prolonged exposure. When the *C*_m_ values reach the clinical MEC, inhibition of glucose transport would be manifested. Importantly, the once accumulated alcohols would not be almost redistributed back into the water because of their high P values, thereby unilaterally accumulating in membrane. These effects may contribute to long-term inhibition of glucose transport, resulting in the development of type 2 diabetes, as seen in decreased yeast cell viability.^18^ Although cutoff alcohols are not typically ingested, similar chemicals with long-alkyl chains, such as emulsifiers in foods and related compounds are. Because the mechanism shown in Figure 7 applies to such compounds, persistent and excessive consumption may contribute to diabetes as chemical factors. Recently, various food additive emulsifiers were reported to be correlated with the risk of type 2 diabetes.^50^ This study provides a theoretical basis for this correlation.

### Membrane concentration based on lipid theory provides the basis for correct evaluation of chemical potency

Chemicals possess intrinsic *P* values (*P*_ow_ and *K*_mw_). In aqueous solution containing cells surrounded by membrane, compounds are distributed according to the P values in the *C*_m_, reflecting the cellular concentration via dissolution in the membrane, and in the *C*_w_, reflecting the non-cellular concentration in water. Lipid theory focuses on *C*_m_, whereas protein theory focuses on administered *C*_w_, which is a major difference between these theories. Distinguishing between these theories is crucial because it is directly linked to correct evaluation of chemical potency..

Empirically, because chemical potency is evaluated as 1/*C*_w_, highly lipophilic chemicals with low *C*_w_ values are considered to have high potency and target small amounts of proteins. QSAR models estimate 1/*C*_w_ from the chemical structure.^11-13^ However, the present models cannot explain the principle of the MO correlation, a prototype of QSAR, and its contradictions, such as the cutoff paradox and activity cliff. This is because QSAR equations define logP ambiguously as a non-dimensional parameter for the hydrophobicity of chemicals and ignore the original meaning of *P* as the ratio. In addition, these equations are only empirical models generated by fitting known data and thus have no biological meaning. Nevertheless, the equations have been interpreted to represent chemical-protein interactions since the 1960s, providing a basis for serious confusion: protein theory.

In aqueous solution, chemicals exist as a mixture of effective and ineffective structures; thus, potency cannot be correctly evaluated without distinguishing these structures. For non-electrolyte alcohols, hydrated monomers formed at a concentration lower than the *S*_w_ are effective structures that dissolve in the membrane and exert effects. However, aggregated droplets formed hydrophobically at a concentration higher than the *S*_w_ are ineffective structure that does not dissolve in it, which is the cause for cutoff paradox.^17,18^ Even at concentrations less than *S*_w_,, antimalarial QC and chloroquine exist as a mixture of its CAD and hydrophilic (HP) forms at physiological pH. CAD is an effective structure that dissolves in the membrane and exerts its effects, whereas the HP is ineffective structure that does not. When CADs are altered to their HP forms through protonation with a slight decrease in pH, the drugs lose their biological activity, which is the cause for resistance.^21^ Thus, it is necessary to measure Cm, which reflects the effective structure, to accurately evaluate chemical potency.

However, estimating *C*_m_ is difficult because the *K*_mw_ values for most chemicals are unknown. Rather than using *C*_m_, *C*_o_ is a useful approximation because the *P*_ow_ and *S*_w_ values for various chemicals can be estimated using QSPR software (Table 1), although corrections may be required for the accurate estimation of highly lipophilic chemicals. When evaluated using 1/*C*_w_, alcohol potency appears to increase exponentially with chain length to inhibit glucose transport. QC was 3-fold less potent than CPZ at pH 7.5, whereas based on 1/*C*_o_, alcohol potency was very low and remained constant regardless of chain length. The potency of QC was 8-fold higher than that of CPZ (Table 1). In addition, although the *C*_w_ cannot be estimated for cutoff alcohols, *C*_o_ can be estimated as the product of *P*_ow_ with *S*_w_, the maximum concentration of hydrated monomer able to dissolve in the membrane.^18^ Presumably, *C*_o_ would have a low value that does not reach a constant value exerting an effect, thereby accounting for the cutoff paradox and drug resistance. Thus, the *C*_o_ reflecting the cellular concentration more accurately evaluates chemical potency than *C*_w_.

## CONCLUSION

*C*_w_, which reflects the non-cellular concentration, cannot accurately evaluate chemical potency and has contributed to long-standing misconceptions, such as the cutoff paradox, drug resistance, and protein theory, and has hindered theoretical drug design. Reconsidering the meaning of great insight for *C* _m_ ^14^ reflecting cellular concentration and estimating *C*_m_ using the present QSPR technology^30^ would correctly reflect potency, thereby providing solutions.

## EXPERIMENTAL SECTION

### Strains, media, plasmids, and growth conditions

Standard genetic manipulation of yeast and DNA manipulation were performed as described previously.^51^ The medium used for the budding yeast *Saccharomyces cerevisiae* was YPD (1% yeast extract, 2% polypeptone, and 2% glucose). The yeast strain used was BY4741 as the wild-type (MATa *his3Δ1 leu2Δ0 ura3Δ0 met15Δ0*) and EBY.S7, deleting all 18 hexose transporter genes (MATa *hxt1-17Δ gal2Δ agt1Δ stl1Δ leu2-3, 112 ura3-52 trp1-289 his3-Δ1 MAL2-8c SUC2 fgy1-1*).^23^ EBY.S7 harboring Hxt2mnx-pVT, the *ADH1* promoter–driven *HXT2* gene on multicopy vector pVT102-U (*2µ ori, URA3*), was used as the HXT2m strain expressing a high affinity hexose transporter alone.^27,52^ Cellular turbidity (OD_600_) was monitored by a UV spectrophotometer (Ultrospec 1100 Pro, GE Healthcare, USA), and the cell numbers were counted with hematocytometer. The values were determined in YPD cultures of HXT2m (OD_600_ = 1: 1.2 × 10^7^ cells/ml).

### Chemicals

*n-*Alcohols (C_2_−C_14_), methylene blue (MB) tetrahydrate, and trisodium citrate dihydrate were purchased from Fujifilm Wako Pure Chemical Corporation (Tokyo, Japan). All compounds are >95% pure by HPLC. Physicochemical properties of the Alcohols were obtained from the PubChem database (https://pubchem.ncbi.nlm.nih.gov/).

### Determination of the uptake rate of 2DG and kinetic analysis

The initial rate of hexose uptake was determined under zero-trans entry conditions with 2-deoxyglucose (2DG) as substrate using the bioluminescent assay, which is based on measuring 2DG6P converted from 2DG transported via a hexose transporter (Glucose uptake-Glo™ assay, Promega). Previous method^21^ was improved for kinetic assay of yeast Hxt2, a high affinity hexose transporter, as follows. Original reductase solution by manufacturer was diluted 2-fold, and the reductase substrate solution was diluted 10-fold to get a linear correlation between low level of 2DG and amount of luminescence. Logarithmic cultures of the HXT2m strain grown in YPD were washed twice with sodium phosphate buffer (PB: 0.1 M, pH 7.4) and suspended in the same buffer as the glucose-starved cells. Eighty microliters of the PB solution containing the indicated concentrations of the alcohols (C_2_, C_4_, C_6,_ and C_8_) were mixed with 10 µl of the above cell suspension (2.4 × 10^7^ cells) at 25°C for 30 min. Then, 10 µl of 2DG was sequentially mixed with the alcohols-treated cells at the indicated concentrations (100 µl total, 2.4 × 10^6^ cells) and incubated for 10 seconds at 25°C. The reaction mixtures were snap-frozen in liquid nitrogen. The mixtures were dissolved on ice and centrifuged at 4°C, and the cell pellets were mixed with 44 µl of 1.75M HClO_4_ (10 %) for 10 min on ice. The acid extracts containing cells were neutralized with 6 µl of 5 M K_2_CO_3_ for 10 min on ice. After centrifugation, the supernatants of the neutralized solutions containing 2DG6P were diluted 20-fold in sodium phosphate buffer (0.1 M, pH 7.4), and the salt pellets were dissolved in 1 ml of water to measure the OD_600_. Tree microliters of the diluted solution were mixed with equivalent amounts of the 2DG6P detection reagent and incubated at 25°C for 30 min, and then the amounts of 2DG6P were determined using a luminometer (AB-120, ATTO, Tokyo, Japan) according to the manufacturer’s instructions. The amount of 2DG6P arising from the transported 2DG was normalized by subtracting the background luminescence arising from authentic G6P in the yeast cells untreated with 2DG. The initial rate of 2DG uptake was expressed as the unit per cell (nmol/min·OD_600_). For the half inhibitory concentrations (IC_50_), the initial rates were determined at the alcohols concentration in the presence of 1 mM 2DG. For kinetic parameters (*K*_m_ and *V*_max_), the initial rates were determined in the presence of 2DG (0–4 mM) and the indicated alcohols at around their IC_50_ values. It should be noted that this method was not applicable for the low affinity hexose transporter, possibly because a high concentration of transported 2DG inhibited hexokinase activity.

### Determination of drug efflux activity by spectroscopic method.^28^

The HXT2m cells were prepared by the manner as 2DG uptake. One hundred microliters of PB solution containing 2.4 × 10^6^ cells were treated with the indicated concentrations of the above alcohols (C_2_−C_8_) at 25°C for 30 min. The reaction mixtures were centrifuged, and the cell pellets were washed once with 100 µl of 0.5% sodium citrate. The pellets were mixed with 100 µl of MB solution containing 0.5% sodium citrate and 0.0025% of methylene blue for 2 min and centrifuged. Eighty microliters of the supernatants were measured on 96 wells plate at 665 nm using a microplate reader (MTP-300, Corona Electric, Ibaragi, Japan). The change of MB absorbance in the supernatants is affected by initial concentration of MB solution; thus, the absorbance was standardized using the following equation:

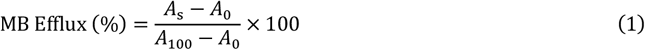

where As is the absorbance of the supernatant of the sample, and *A*_100_ and *A*_0_ are the absorbances of the supernatants for the same cell number of 100% viable and 100% dead cells, respectively. The 100% dead cells were prepared by treatment with 70% ethanol for 30 min. When *A*_s_ was greater than *A*_100_, the viability was considered as 100%, and when smaller than A0, considered as 0%. This MB efflux highly correlates with cell viability.^28^

### Determination of intracellular ATP level

Intracellular ATP levels of yeast were measured in the presence of glucose.^29^ The HXT2m strain were grown in YPD buffered with PB (0.1 M, pH 7.4), and 100 µl of the logarithmic cultures containing 2.4 × 10^6^ cells were treated with the indicated alcohols at 25°C for 30 min. The cell suspensions were centrifuged, and the pellets were mixed with 10 µl of 10% TCA and then diluted with 990 µl of ATP buffer (25mM Tris-HCl pH7.5, 2mM EDTA, final 0.1% TCA). Five microliters of the diluted cell suspensions were mixed with equivalent amounts of the luciferase/luciferin mixture from the ATP extraction system kit (LL-100-1; Toyo Ink, Tokyo, Japan). The amounts of ATP were measured using the luminometer according to the manufacturer’s instructions. The intracellular ATP level extracted by TCA does not reflect absolute levels because of low extraction efficiency. Thus, the decrease in ATP by alcohols was calculated as percentages to the intracellular ATP level in the cell untreated with alcohol.

### Determination of the Minimum Inhibitory Concentration

The minimum inhibitory concentrations (MICs) for the yeast growth were determined as described previously.^17^ One hundred microliters of logarithmic BY4741 and HXT2m cells in YPD (10 mM PB, pH7.4) containing 6.5 × 10^4^ cells or 6.5 × 10^2^ (1/100) with 2% or 8% glucose were dispensed into 96-well microtiter plates (Iwaki, Tokyo, Japan) and incubated statically at 25 °C. The same amounts of cells containing C_2_ (10 M) and C_4_ (1M) were added directly to the above cell suspensions, and 2-fold serial dilutions were performed in the wells. For the alcohols (C_6_-C_14_), 2-fold serial dilutions were performed in DMSO and 1 μL of the alcohols were dispensed into wells before adding the above cell cultures. Because the doubling times for BY4741 and HXT2m strains in YPD containing 2% glucose were 1.81 and 3.06 hours, these cells were incubated for 3 and 5 days, respectively, for reaching same numbers of division. The MIC was the lowest concentration that completely inhibited microbial growth, as judged by visual observation.

### Computational methods

Distribution coefficients (log*D*) of the alcohols, quinacrine (QC), and chlorpromazine (CPZ) at pH7.4 were estimated using MarvinSketch software package (version 23.16) (ChemAxon, Hungary) (accessed October, 2025). The consensus log*P* method was applied to calculate the distribution coefficient of each molecule.

### Data analysis

The IC_50_, *K*_m_, and *V*_max_ values were determined by the nonlinear least-squares method of the log-logistic^53^ and Michaelis-Menten^54^ models using the Solver add-in bundled with Microsoft Excel. Statistical analyses were performed by unpaired Student’s *t*-test and one-way analysis of variance with the Tukey-Kramer multiple comparison test, using the R statistical software package (ver. 3.5.1; R Development Core Team 2018).

## AUTHOR INFORMATION

### Notes

The authors declare no competing financial interest.

## ACKNOWLEDGMENT

We thank Drs. Y. Inoue, W. Nomura, and K. Yamada of Kyoto University, T. Kasahara of Teikyo University, and Y. Osakabe of Institute of Science Tokyo for providing plasmids and strains.

## ABBREVIATIONS

HXT: hexose transporter
MO: Meyer-Overton
C_2_: ethanol
C_4_: 1-butanol
C_6_: 1-hexanol
C_8_: 1-octanol
C_10_: 1-decanol
C_12_: 1-dodecanol
C_14_: 1-tetradecanol
QSAR: Quantitative Structure–Activity Relationship
QSPR: Quantitative Structure–Property Relationship
*S*_w_: saturated aqueous solubility
IC_50_: half inhibitory concentration
MEC: minimum effective concentrations
2DG: 2-deoxyglucose
MB: methylene blue
MIC: minimum inhibitory concentration
logP: logarithm of partition coefficient
logD: logarithm of distribution coefficient
CAD: cationic amphiphilic drug
HP: hydrophilic
OD: optical density
QC: quinacrine
CQ: chloroquine
CPZ: chlorpromazine
YPD: yeast extract,polypeptone,dextrose.

## REFERENCES

(1) Meyer, H. Zur theorie der Alkoholnarkose. Arch. Exp. Pathol. Pharmakol. 1899, 42, 109–118.

(2) Overton, C. E. Über die osmotischen Eigenschaften der lebenden Pflanzen und Tierzelle. Vierteljahresschrift Naturforsch. Ges. Zürich 1895, 40, 159–201.

(3) Pringle, M. J. Brown, K. B. Miller, K. W. Can the lipid theories of anesthesia account for the cutoff in anesthetic potency in homologous series of alcohols? Mol. Pharmacol. 1981, 19, 49–55.

(4) Antkowiak, B. How do general anaesthetics work? Naturwissenschaften 2001, 88, 201–213.

(5) Gill, C. O. Ratledge, C. Toxicity of n-alkanes, n-Alk-1-enes, n-alkan-1-ols and n-alkyl-1-bromides towards yeasts. Microbiology 1972, 72, 165–172.

(6) Teh, J. S. Toxicity of short-chain fatty acids and alcohols towards Cladosporium resinae. Appl. Microbiol. 1974, 28, 840–844.

(7) Kubo, I. Muroi, H. Kubo, A. Structural functions of antimicrobial long-chain alcohols and phenols. Bioorg. Med. Chem. 1995, 3, 873–880.

(8) Uesono, Y. Toh-E, A. Kikuchi, Y. Terashima, I. Structural analysis of compounds with actions similar to local anesthetics and antipsychotic phenothiazines in yeast. Yeast 2011, 28, 391–404.

(9) Roth, S. Seeman, P. The membrane concentrations of neutral and positive anesthetics (alcohols, chlorpromazine, morphine) fit the Meyer-Overton rule of anesthesia; negative narcotics do not. Biochim. Biophys. Acta. 1972 255, 207–219.

(10) Ingólfsson, H. I. Andersen, O. S. Alcohol’s effects on lipid bilayer properties. Biophys. J. 2011, 101, 847–855.

(11) Hansch, C. Maloney, P. P. Fujita, T. Muir, R. M. Correlation of biological activity of phenoxyacetic acids with Hammett substituent constants and partition coefficients Nature 1962, 194, 178–180.

(12) Hansch, C. Fujita, T. p-σ-π Analysis. A Method for the Correlation of Biological Activity and Chemical Structure. J. Am. Chem. Soc. 1964, 86, 1616–1626.

(13) Hansch, C. Quantitative approach to biochemical structure-activity relationships. Acc Chem. Res. 1969, 2, 232–239.

(14) Meyer, K. H. Hemmi, H. Beitrage zur Theorie der Narkose, III. Biochem. Z 1935, 277, 39–71.

(15) Franks, N. P. Lieb, W. R. Molecular and cellular mechanisms of general anaesthesia. Nature 1994, 367, 607–614.

(16) Franks, N. P. Lieb, W. R. Mapping of general anaesthetic target sites provides a molecular basis for cutoff effects. Nature 1985, 316, 349–351.

(17) Matsumoto, A. Uesono, Y. Physicochemical Solubility of and Biological Sensitivity to Long-Chain Alcohols Determine the Cutoff Chain Length in Biological Activity. Mol. Pharmacol. 2018, 94, 1312–1320.

(18) Matsumoto, A. Adachi, H. Terashima, I. Uesono, Y. Escaping from the Cutoff Paradox by Accumulating Long-Chain Alcohols in the Cell Membrane. J. Med. Chem. 2022, 65, 10471–10480.

(19) Matsumoto, A. Uesono, Y. Establishment of the Meyer-Overton correlation in an artificial membrane without protein. Biochim. Biophys. Acta Gen. Subj. 2024, 1868, 130717.

(20) Uesono, Y. Toh-e, A. Kikuchi, Y. Araki, T. Hachiya, T. Watanabe, CK. Noguchi, K. Terashima, I. Local Anesthetics and Antipsychotic Phenothiazines Interact Nonspecifically with Membranes and Inhibit Hexose Transporters in Yeast. Genetics 2016, 202, 997–1012.

(21) Kitagawa, T. Matsumoto, A. Terashima, I. Uesono, Y. Antimalarial Quinacrine and Chloroquine Lose Their Activity by Decreasing Cationic Amphiphilic Structure with a Slight Decrease in pH. J. Med. Chem. 2021, 64, 3885–3896.

(22) Drew, D. North, R.A. Nagarathinamm, K. Tanabe, M. Structures and General Transport Mechanisms by the Major Facilitator Superfamily (MFS). Chem. Rev. 2021, 121, 5289–5335.

(23) Wieczorke, R. Dlugai, S. Krampe, S. Boles, E. Characterisation of mammalian GLUT glucose transporters in a heterologous yeast expression system. Cell. Physiol. Biochem. 2003, 13, 123–134.

(24) Ozcan, S. Johnston, M. Function and regulation of yeast hexose transporters. Microbiol. Mol. Biol. Rev. 1999, 63, 554–569.

(25) Alifimoff, J. K. Firestone, L. L. Miller, K. W. Anaesthetic potencies of primary alkanols: implications for the molecular dimensions of the anaesthetic site. Br. J. Pharmacol. 1989, 96, 9–16.

(26) Reifenberger, E. Boles, E. Ciriacy, M. Kinetic Characterization of individual hexose transporters of Saccharomyces cerevisiae and their relation to the triggering mechanisms of glucose repression. Eur. J. Biochem. 1997, 245, 324–333.

(27) Kasahara, T. Kasahara, M. Transmembrane segments 1, 5, 7 and 8 are required for high-affinity glucose transport by Saccharomyces cerevisiae Hxt2 transporter. Biochem. J. 2003, 372, 247–252.

(28) Matsumoto, A. Terashima, I. Uesono, Y. A rapid and simple spectroscopic method for the determination of yeast cell viability using methylene blue. Yeast. 2022, 39, 607–616.

(29) Uesono, Y. Ashe, M. P. Toh-E, A. Simultaneous yet independent regulation of actin cytoskeletal organization and translation initiation by glucose in Saccharomyces cerevisiae. Mol. Biol. Cell. 2004, 15, 1544–1556.

(30) Leo, A. J. Calculating log Poct from structures. Chemical reviews. 1993, 93, 1281–1308.

(31) Zhao, J. Wu, J. Heberle, F. A. Mills, T. T. Klawitter, P. Huang, G. Costanza, G. Feigenson, G. W. Phase studies of model biomembranes: complex behavior of DSPC/DOPC/cholesterol, Biochim. Biophys. Acta Biomembr. 2007, 1768, 2764–2776.

(32) Pike, L. J. Rafts defined: a report on the keystone symposium on lipid rafts and cell function, J. Lipid Res. 2006, 47, 1597–1598.

(33) Yoshida, A. Wei, D. Nomura, W. Izawa, S. Inoue, Y. Reduction of glucose uptake through inhibition of hexose transporters and enhancement of their endocytosis by methylglyoxal in Saccharomyces cerevisiae. J. Biol. Chem. 2012, 287, 701–711.

(34) Janoff, A. S. Pringle, M. J. Miller, K. W. Correlation of general anesthetic potency with solubility in membranes. Biochim Biophys Acta. 1981, 649, 125–128.

(35) Araki, T. Toh-e, A. Kikuchi, Y. Watanabe, C. K. Hachiya, T. Noguchi, K,; Terashima, I. Uesono, Y. Tetracaine, a local anesthetic, preferentially induces translational inhibition with processing body formation rather than phosphorylation of eIF2α in yeast. Curr. Genet. 2015, 61, 43–53.

(36) van der Rest, M. E. Kamminga, A. H. Nakano, A. Anraku, Y. Poolman, B. Konings, W. N. The plasma membrane of Saccharomyces cerevisiae: structure, function, and biogenesis. Microbiol. Rev. 1995, 59, 304–322.

(37) Sonner J. M., Cantor, R. S. Molecular mechanisms of drug action: an emerging view. Annu. Rev. Biophys. 2013, 42, 143–167.

(38) Payandeh, J. Volgraf, M. Ligand binding at the protein-lipid interface: strategic considerations for drug design. Nat. Rev. Drug Discov. 2021 20, 710–722.

(39) Franks, N. P. Lieb, W. R. Do general anaesthetics act by competitive binding to specific receptors? Nature 1984, 310, 599–601.

(40) Suzuki, H., Kawarabayasi, Y. Kondo, J. Abe, T. Nishikawa, K. Kimura, S. Hashimoto, T. Yamamoto, T. Structure and regulation of rat long-chain acyl-CoA synthetase. J Biol Chem 1990, 265, 8681–8685.

(41) Mofford, D. M. Liebmann, K. L. Sankaran Kiran Kumar Reddy, G. S. Randheer Reddy G. Miller, S. C. Luciferase Activity of Insect Fatty Acyl-CoA Synthetases with Synthetic Luciferins. ACS Chem. Biol. 2017 12, 2946–2951.

(42) Oba, Y. Ojika, M. Inouye, S. Firefly luciferase is a bifunctional enzyme: ATP-dependent monooxygenase and a long chain fatty acyl-CoA synthetase. FEBS Lett. 2003 540, 251–254.

(43) Hisanaga, Y. Ago, H. Nakagawa, N. Hamada, K. Ida, K. Yamamoto, M. Hori, T. Arii, Y. Sugahara, M. Kuramitsu, S. Yokoyama, S. Miyano, M. Structural basis of the substrate-specific two-step catalysis of long chain fatty acyl-CoA synthetase dimer. J. Biol. Chem. 2004, 279, 31717–31726.

(44) Alkire, M. T. Haier, R. J. Shah, N. K. Anderson, C. T. Positron emission tomography study of regional cerebral metabolism in humans during isoflurane anesthesia. Anesthesiology 1997, 86, 549–557.

(45) Languren, G. Montiel, T. ; Julio-Amilpas, A. Massieu, L. Neuronal damage and cognitive impairment associated with hypoglycemia: an integrated view. Neurochem. Int. 2013 63, 331–343.

(46) Nakamura, Y. Matsumura, K. Miura, K. Kurokawa, H. Abe I. Takata, Y. Cardiovascular and sympathetic responses to dental surgery with local anesthesia. Hypertens. Res. 2001, 24, 209–214.

(47) Haupt, D. W. Newcomer, J. W. Hyperglycemia and antipsychotic medications. J. Clin. Psychiatry 2001, 27, 15–26.

(48) Onishi, Y. Honda, M. Ogihara, T. Sakoda, H. Anai, M. Fujishiro, M. Ono, H. Shojima, N. Fukushima, Y. Inukai, K. Katagiri, H. Kikuchi, M. Oka, Y. Asano, T. Ethanol feeding induces insulin resistance with enhanced PI 3-kinase activation. Biochem. Biophys. Res. Commun. 2003, 303, 788–794.

(49) Wickramasinghe, P. B. Qian, S. Langley, L. E. Liu, C. Jia, L. Hepatocyte Toll-Like Receptor 4 Mediates Alcohol-Induced Insulin Resistance in Mice. Biomolecules 2023, 13:454.

(50) Salame, C. Javaux, G. Sellem, L. Viennois, E. Szabo de Edelenyi, F. Agaësse, C. De Sa, A. Huybrechts, I. Pierre, F. Coumoul, X. Julia, C. Kesse-Guyot, E. Allès, B. Fezeu, L. K. Hercberg, S. Deschasaux-Tanguy, M. Cosson, E. Tatulashvili, S. Chassaing, B. Srour, B. Touvier, M. Food additive emulsifiers and the risk of type 2 diabetes: analysis of data from the NutriNet-Santé prospective cohort study. Lancet Diabetes Endocrinol. 2024, 12, 339–349.

(51) Guthrie, C. Fink, G. R., Eds. Guide to Yeast Genetics and Molecular Biology, 194. Academic Press/Elsevier: Cambridge, 1991.

(52) Yoshida, A. Wei, D. Nomura, W. Izawa, S. Inoue Y. Reduction of glucose uptake through inhibition of hexose transporters and enhancement of their endocytosis by methylglyoxal in Saccharomyces cerevisiae. J. Biol. Chem. 2012, 287, 701–711.

(53) Onofri, A. BIOASSAY97: a new EXCEL VBA macro to perform statistical analyses on herbicide dose–response data. Riv. Ital. Agrometeorol. 2005, 3, 40–45.

(54) Kemmer, G, Keller, S. Nonlinear least-squares data fitting in Excel spreadsheets. Nat. Protoc. 2010, 5, 267–81.

